# Self-supervised vision transformers accurately decode cellular state heterogeneity

**DOI:** 10.1101/2023.01.16.524226

**Authors:** Ramon Pfaendler, Jacob Hanimann, Sohyon Lee, Berend Snijder

## Abstract

Characterising cellular phenotypic heterogeneity is essential to understand the relationship between the molecular and morphological determinants of cellular state. Here we report that publicly available self-supervised vision transformers (ss-ViTs) accurately elucidate phenotypic stem cell heterogeneity out-of-the-box. Moreover, we introduce scDINO, an adapted ss-ViT trained on five-channel automated microscopy data, attaining excellent performance in delineating peripheral blood immune cell identity. Thus, ss-ViTs represent a leap forward in the unsupervised analysis of phenotypic heterogeneity.

## Main

Advances in microscopy-based high-content imaging have revolutionised the phenotypic characterisation of single cells in their microenvironment^1,2^. Classically, conventional image analysis quantifies the observed phenotypes by predefined features, which describe important and interpretable cellular traits including size, shape, and texture, enabling cellular phenotype classification^3–5^. However, contemporary advances in supervised deep learning (DL), particularly convolutional neural networks (CNN), have shown superior classification performance compared to conventional image analysis based on improved extraction of image-intrinsic features^6,7^. This has led to an unprecedented view on the relationship between cellular morphology and cellular state^8–12^. The performance of CNNs has recently been surpassed by the introduction of Vision Transformers (ViTs)^13^. While CNNs analyse input images explicitly as pixel arrays, ViTs divide an image into fixed-size patches with positional embeddings and implement so-called self-attention mechanisms, which enhances the representation of spatial relationships between distant image locations^13–15^. Self-attention can be visualised to reveal what the model focuses on. Furthermore, ViTs implement a class (CLS) token that represents the global image information, enabling accurate clustering of images by their similarity^13^. These clustering accuracies are commonly assessed by measuring if the k-nearest neighbours (k-NN) of each image contain the same label.

Recently, DL is seeing a shift from supervised learning towards fully self-supervised learning^16^. With self-supervision, models learn meaningful features from the data itself, without the need for manual data labelling, thereby limiting bias. In a seminal biological study, self-supervised DL was successfully applied to profile subcellular protein localisation in microscopy data using an autoencoder^17^. However, as the authors point out, additional work is needed to disentangle cellular phenotypic heterogeneity by self-supervised deep learning^17^.

One self-supervised training strategy, coined DINO, was recently used to train self-supervised ViTs (ss-ViTs) leading to excellent k-NN classification performance^18^. Here, a student network is trained to match the internal representation (or probability distribution) of a teacher network on differently augmented versions of the same image (Fig. 1A). DINO-trained ss-ViTs learn features containing explicit information about the object boundaries in RGB-based images and accurately cluster similar image subjects based on their CLS token output^18^. Recent studies have demonstrated that DINO-based models can learn to pay attention to population-level tissue phenotypes^19,20^. However, ss-ViTs are yet to be used for the study of cellular state heterogeneity on a single-cell level.

**Figure 1:**
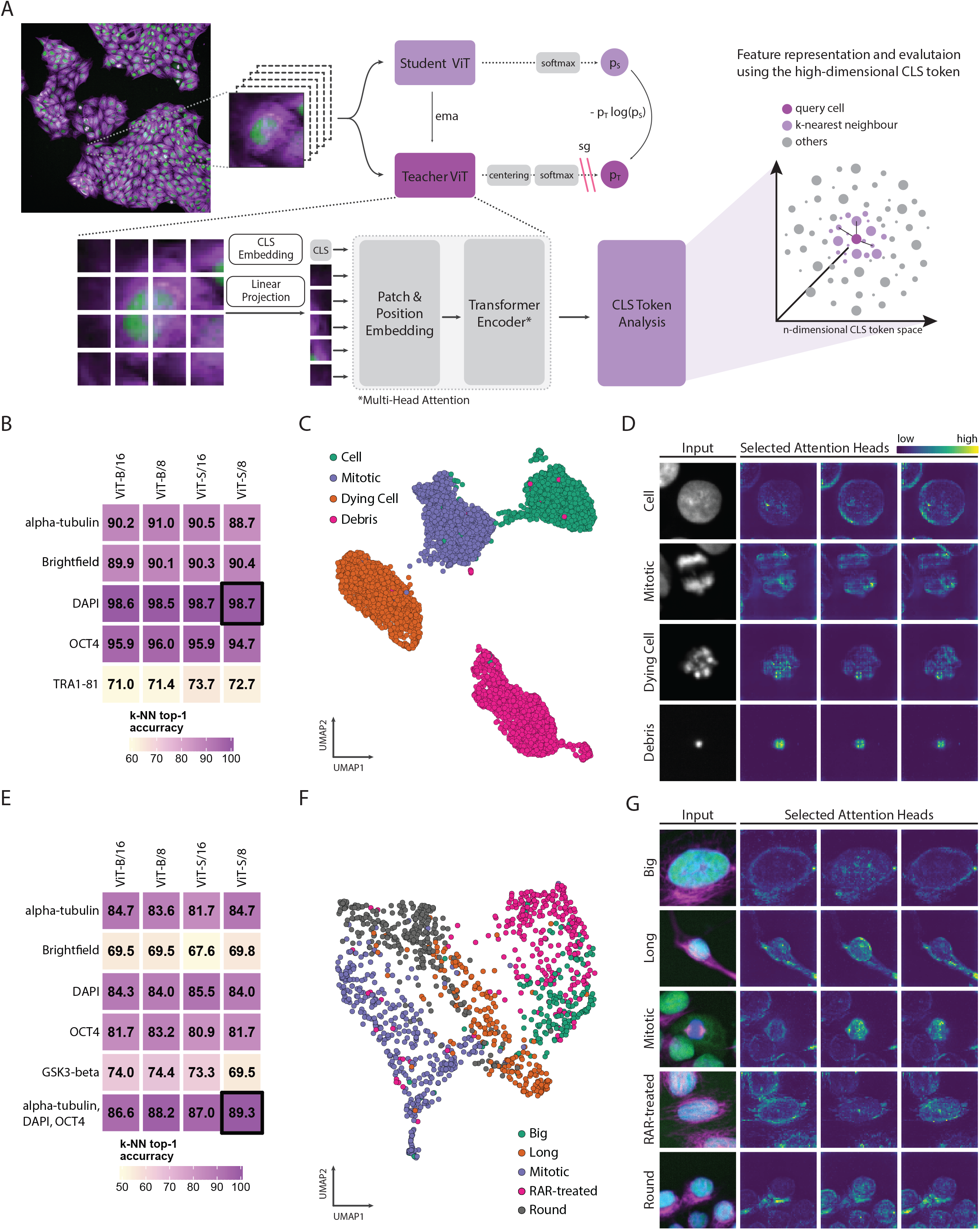
ss-ViTs decipher cellular state heterogeneity in human stem cells. **A,** Workflow for single-cell phenotyping with DINO-based self-supervised vision transformers (ss-ViTs). Upper part: DINO-based self-supervised learning approach incorporating a student and teacher ViT. Lower part: ViT-based image embedding and feature representations (CLS token) used for single-cell clustering. **B,** Heatmap displaying k-NN top-1 accuracies of microscopy-adjusted ImageNet-trained ss-ViTs per channel (rows) and architectures (columns) for the four-class iPSC dataset (Cell, Mitotic, Dying Cell, Debris). Black-outlined model is used for further analysis. **C,** UMAP projection of the CLS tokens of 5,922 cells evaluated using the highlighted model. Each dot corresponds to a single cell, colour-coded by their label. **D,** Representative images of each class with one greyscale channel (DAPI), followed by visualisation of selected self-attention heads. Each image crop measures 30μm x 30μm. **E-G,** as in **B-D**, but for the analysis of iPSC morphology across three channels (α-tubulin, magenta; DAPI, blue; OCT4, green). Each image crop measures 45μm x 45μm. **F,** UMAP shows 1312 cells.

Here, we evaluate the use of DINO-trained ss-ViTs for the analysis of single-cell phenotypic heterogeneity captured by high-throughput automated microscopy. We first assessed four previously reported ss-ViTs^18^ that had been DINO-trained on a publicly available dataset of RGB-based images capturing everyday subjects (ImageNet^21^), yet differed in network configuration (‘big’ and ‘small’) and patch size (8×8 or 16×16 pixels). Contrary to the ImageNet dataset, in fluorescence microscopy, variable numbers of channels assess biological properties individually, and the order of channels in the final image is arbitrary (i.e. sequence invariant). To make the four pretrained ss-ViTs applicable to microscopy images, we therefore adapted the models to accept variable numbers of input channels, and averaged the RGB-specific embedding weights, giving each microscopy channel an equal weight.

As a proof-of-principle, we applied the adjusted models first to a relatively simple biological problem: the separation of healthy, dividing, and dying cells, as well as debris, in five-channel high-content imaging data of human induced pluripotent stem cells (iPSCs). We created nucleus-centred single-cell sub-images of 5,922 iPSCs, which we manually split into the four classes, and made publicly available for reproducibility^22^. In addition to label-free bright-field imaging, the four fluorescent channels recorded different markers, specifically DNA staining (DAPI), stem cell marker expression (OCT4 and TRA-1-81), and cytoskeletal properties (α-tubulin). We then analysed each channel individually with the four adjusted ss-ViTs and evaluated the k-NN accuracies of the CLS token-based image clustering. These accuracies were consistent across the four architectures, yet highly dependent on the analysed image channel. Clustering based on channels capturing nuclear characteristics, i.e. DAPI and OCT4, performed particularly well. Using the DAPI channel, all four ss-ViTs achieved k-NN top-1 accuracies above 98%, showing excellent clustering of the four classes (Fig. 1B). UMAP^23^ embedding all cells based on their DAPI-specific CLS tokens from ViT-S/8 (i.e. the small network configuration with patch size of 8×8 pixels) showed a clear separation of cellular states, demonstrating that the models recognised class-specific morphological features without any supervision (Fig. 1C). Furthermore, assessing the output of the self-attention mechanism showed that the model distinguished characteristic nuclear properties in the respective classes (Fig. 1D). Summarised, we find that ImageNet-pretrained ss-ViTs recognise morphological characteristics, leading to remarkably accurate clustering of simple cellular phenotypes, without any training on microscopy images.

Next, we explored if ss-ViTs can capture more subtle morphological differences among healthy iPSCs. Given their developmental plasticity, iPSCs are known to generate a broad range of cellular states in response to perturbations. We challenged iPSCs with several small-molecule compounds and stained their resulting nuclear organisation (DAPI), stem cell potential (OCT4), signalling state (GSK3-β), and cytoskeleton (α-tubulin). From this dataset, we collected 1,312 cells approximately equally split across five visually-identified morphological classes, referred to as ‘Long’, ‘Round’, ‘Big’, ‘Mitotic’, and a perturbation-induced phenotype labelled ‘RAR-treated’, also available in the public resource^22^. Analysing all single-channels, the four ss-ViT models discerned the single-cell phenotypes with good accuracies based on either DAPI or α-tubulin staining (83.2% and 84.7% respectively; Fig. 1E). However, analysis of three-channel images, combining α-tubulin, DAPI, and OCT4, increased the k-NN top-1 accuracy to 89.3%, illustrating how additional biological information was beneficial to distinguish these five morphological classes (Fig. 1E). Projecting the CLS tokens of the 1,312 iPSCs evaluated with ViT-S/8 using UMAP demonstrated that this model recognised subtle phenotypic differences and preserved an intuitive global clustering across the five morphologies (Fig. 1F). This was further reflected in the self-attention maps, which revealed that the model perceived class-specific subcellular features, particularly focussing on the nuclear envelope and the nucleus-peripheral cytoskeleton (Fig. 1G).

We wondered if DINO-trained ss-ViTs can decipher the cellular state heterogeneity and multilineage identities of human peripheral blood mononuclear cells (PBMCs). We made use of our previously reported, publicly available high-throughput imaging dataset (‘Phenoplex‘^24^), containing 89,564 training and 24,000 test images of eight different manually-annotated immune cell classes from multiple healthy donors^12^. In this dataset, all five channels combined with morphological information were previously necessary to discriminate the eight different classes by supervised deep learning, as lineage-specific markers were multiplexed in unique combinations across the fluorescent channels^12^. Therefore, we evaluated the performance of the ImageNet-pretrained ss-ViTs on five-channel images of a subset of 8,000 class-balanced immune cells from the test set. The four models partially captured the eight immune cell classes, with k-NN accuracies around 53% (Fig. 2A). In an attempt to improve performance, we adjusted the training workflow to accept five-channel input images and DINO-trained the ViT-S/16 architecture on the 89,564 Phenoplex training images. We refer to this self-supervised instance as ‘single-cell DINO’ (scDINO), available online^25^. scDINO outperformed the ImageNet-trained ss-ViTs already after a few training epochs, reaching a remarkable k-NN accuracy of 94.6% on this eight-class problem after 100 epochs (Fig. 2B). Visualisation of the CLS token space of the 8,000 test cells by UMAP showed scDINO’s capabilities in deciphering subtle differences in immune cell phenotypes (Fig. 2C). It recognised the importance of both lineage-specific marker expression and cellular morphology as outlined in the attention maps of the eight different immune cell types. This was reflected by the strong UMAP separation of T cells and monocytes, whose markers were multiplexed in the same channels, thereby requiring attention to cell morphology for their separation (Fig. 2C).

**Figure 2:**
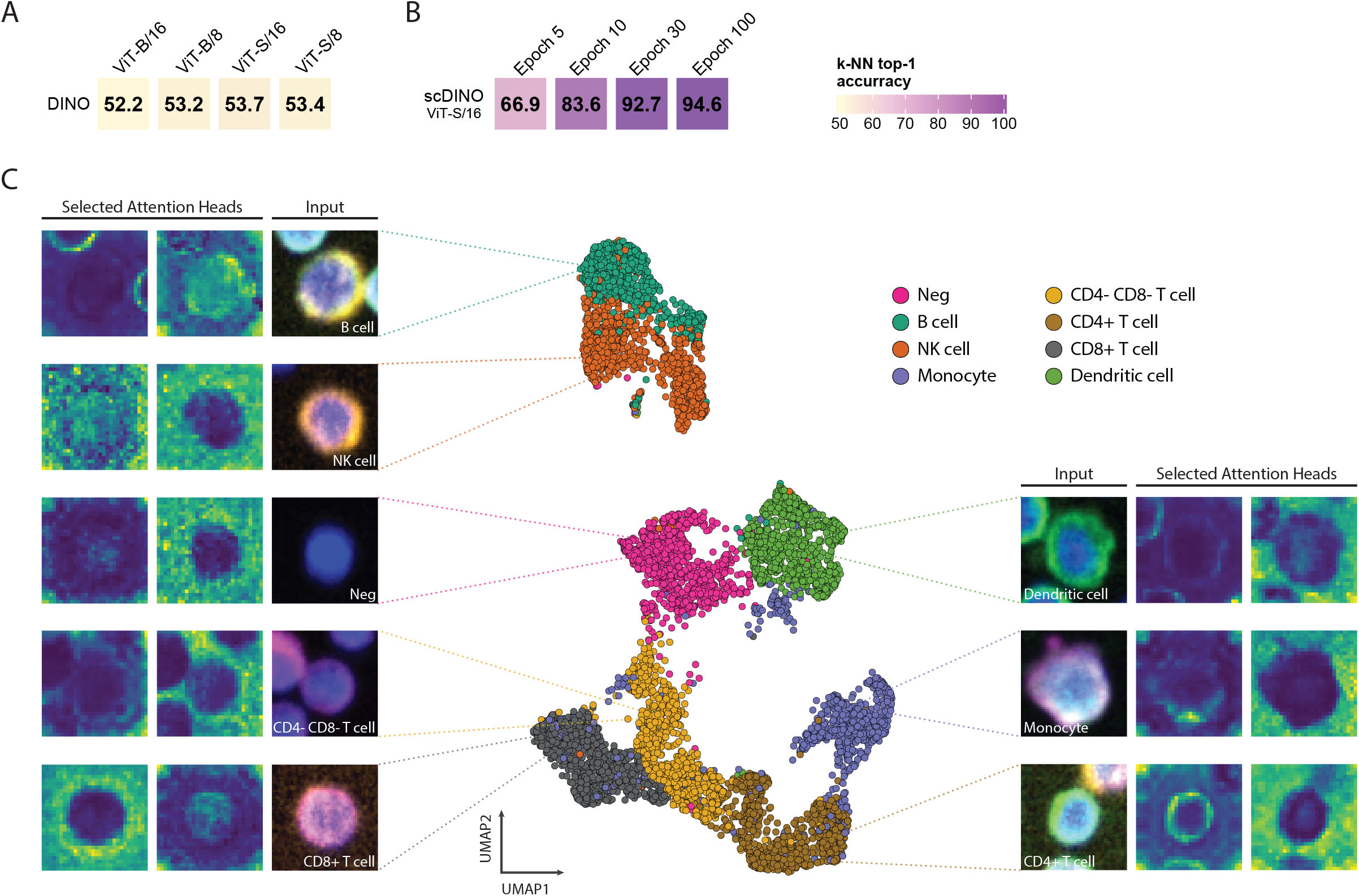
scDINO decodes lineage identity of primary human immune cells. **A,** Heatmap of k-NN top-1 accuracies of different ImageNet-pretrained ss-ViTs (columns are architectures). **B, as in A,** but for scDINO (columns are training epochs). **C,** UMAP of the CLS tokens of 8,000 immune cells (n=100 per donor and class) evaluated with scDINO trained for 100 epochs. Each dot corresponds to a single cell, colour-coded by their label. Representative (pseudo-RGB) images of each cell class with corresponding visualisation of selected attention heads are shown. Images are connected to their respective cells in the UMAP. Each image crop measures 15μm x 15μm.

Taken together, we illustrate that self-supervised vision transformers recognise subtle cellular phenotypes, capable of decoding cellular state heterogeneity present in primary human cells with remarkable accuracy. We find that publicly available ImageNet-pretrained models can be adapted to the analysis of high-throughput microscopy images. Out-of-the-box, these models are sensitive to cell-specific features, discerning fundamental cellular state differences among iPSCs. Furthermore, we introduce scDINO, an ss-ViT DINO-trained on multiplexed immunofluorescence imaging of primary human immune cells, available online (see Data and Code Availability). scDINO achieves excellent performance in the subpopulation identification of eight human immune cell types, revealing its ability to capture channel-encoded marker and morphological heterogeneity from microscopy images without the need for training labels. We thereby show that ss-ViTs have diverse applications for cell microscopy, ranging from analysis of quality-control, to uncovering phenotypic heterogeneity and lineage identity. Thus, DINO-trained vision transformers represent a significant step forward in the unsupervised discovery and quantification of cellular state heterogeneity in image-based screens.

## Data and Code availability

All data needed to evaluate the conclusions and reproduce the sub-panels of the figures present in this paper are available in the Source Data tables.

The code to run the microscopy-adjusted ImageNet-pretrained models, scDINO training and evaluation is available at https://github.com/JacobHanimann/scDINO.

All training and evaluation image datasets used in this study, as well as scDINO, are available at the FAIR principles^28^-compliant repository of ETH Zurich. Both iPSC single-cell imaging datasets are available at https://doi.org/10.3929/ethz-b-000581447^22^. The Phenoplex immune cell imaging dataset was described previously^12^ and is available at https://doi.org/10.3929/ethz-b-000343106^24^. scDINO trained for 100 epochs on the Phenoplex immune cell dataset is available at https://doi.org/10.3929/ethz-b-000582208^25^.

## Material and Methods

### Ethical statement

Human peripheral blood mononuclear cells (PBMCs) were isolated from buffy coats of coded healthy donors provided by the Blutspende Zürich, under a study protocol approved by the Cantonal Ethics Committee, Zürich (KEK Zürich, BASEC-Nr 2019-01579), as described previously^12^.

### iPSC generation from hematopoietic stem cells

From the extracted PBMCs, we further isolated the CD34+ hematopoietic stem cells (HSCs) using magnetic-activated cell sorting according to the manufacturer’s instructions (CD34 MicroBead Kit, Miltenyi Biotec #130-046-702). Subsequently, iPSCs were reprogrammed from HSCs in line with the manufacturer’s protocol (CytoTune-iPS 2.0 Sendai Reprogramming Kit, Thermo Fisher #A16517). In short, CD34+ cells were transduced with Sendai viruses containing three Yamanaka-factors-containing vectors: polycistronic Klf4–Oct3/4–Sox2, cMyc, and Klf4. The emergent colonies were identified according to their morphology and manually picked. Each of the picked clones was then expanded for cryopreservation. iPSC lines were routinely tested negative for Mycoplasma and their pluripotency confirmed by immunofluorescence (OCT4, Nanog, TRA-1-81, TRA-1-60) as well as differentiation into the three germ layers (STEMdiff Trilineage Differentiation kit, #05230, StemCell Technologies).

### Cell culture of IPSCs

iPSCs were cultivated on hESC-qualified Matrigel coated (Lot 0048006, Cornig) plates according to the manufacturer’s instructions at normoxic (5% CO2) and in humidified conditions at 37°C. iPSCs were grown in mTeSR Plus and fed according to manufacturer’s instructions (StemCell Technologies). For general culture, iPSCs were cultivated in 6- or 12-well plates (Nunc) and split using PBS supplemented with 0.5mM EDTA every 2-4 days. For imaging, iPSCs were dissociated to single-cells using 0.75X TrypLE (Thermo Fisher) and subsequently 1500-2000 cells were plated onto Matrigel-coated 384-well plates (PhenoPlate 384-well, Perkin Elmer, #6057308) with mTeSR Plus supplemented with 10uM Y-27632 (Peprotech, #1293823). After 24h, medium was replaced with mTeSR Plus and cells were cultured until fixation.

### Small molecule perturbation of iPSCs

For small molecule perturbations, iPSCs were seeded as described above. After 24h in mTESR Plus supplemented with 10uM Y-27632 (Peprotech), medium was replaced with mTeSR Plus containing one of the following small molecules (all from Sigma) for 72h: 5uM CHIR99021, 20uM AM580, 20um A83-01, 1uM Lysophosphatidic acid, or 0.4% DMSO (control).

### Immunostaining of iPSCs

Cells were fixed in 4% (w/v) formaldehyde (Sigma-Aldrich) in PBS for 15 min at room temperature (RT). The fixative was subsequently removed and the cells washed with PBS once using a plate washer (HydroSpeed, Tecan), before permeabilisation and nuclear staining using PBS supplemented with 2ug/mL DAPI (Biolegend, #422801, stock solution 10mg/ml), 5%FBS (Gibco, #10500064), and 0.1% Triton-X100 (Sigma-Aldrich, #T8787) for 1 hour at RT in the dark. Prior to immunostaining, cells were washed with PBS twice using the plate washer. All iPSCs were stained in PBS with 5% FBS overnight at 4°C with the following antibodies and dilutions: Alexa Fluor^®^ 647 anti-tubulin α (1:1000, Biolegend, #627908, clone 10D8), and Alexa Fluor ^®^ 488 anti-Oct-4A (1:500, Cell Signaling, #5177S, clone C30A3). Additionally, the four-class iPSC dataset (Cell, Mitotic, Dying cell, Debris) was stained with PE anti-human TRA-1-81 (1:300, Biolegend, #330708, clone TRA-1-81), while the five-class iPSC morphology dataset was stained with Alexa Fluor ^®^ 555 anti-GSK3 beta (1:300, abcam, ab201738, clone Y174). The next day, the staining solution was aspirated, cells washed with PBS once, and then kept in PBS at 4°C until imaging.

### Imaging

Cells were imaged using a PerkinElmer Operetta CLS automated spinning-disk confocal microscope. Each well of a 384-well plate was imaged using a 20x high-NA (0.8) air objective with 5×5 non-overlapping images, covering the whole well surface. The images were taken sequentially from the brightfield (650-760 nm), DAPI/Nuclear signal (435-480 nm), GFP/Green signal (500-550 nm), PE/Orange signal (570-630 nm) and APC/Red signal (650-760 nm) channels. Subsequently, the raw .tiff images were transferred from the microscope for further analysis.

### Dataset description and generation - iPSC quality control data

Single cells were segmented based on their nuclear DAPI signal using CellProfiler (version 2.2.0). Subsequently, each cell was mass-centred and cropped as 100×100 pixel images. These single-cell sub-images were then, based on their nuclear morphology, manually curated and labelled into four classes (Debris - Dying Cell - Mitotic Cell - Cell) using custom code in MATLAB R2020a. The dataset is available on ETH Zurich’s Research Collection (https://doi.org/10.3929/ethz-b-000581447)^22^, and contains 5,922 cells, grouped into 1,366 Cells, 1,461 Dying Cells, 1,620 Mitotic Cells, and 1,478 Debris objects.

### Dataset description and generation - iPSC morphologies

Single cells were segmented as described above. Subsequently, each cell was mass-centred and cropped as 150×150 pixel images. These single-cell sub-images were then manually curated and labelled into 5 classes (Long - Round - Mitotic - Big - RAR-treated), according to their combined nuclear and cytoskeletal morphology using custom code in MATLAB R2020a. The dataset is available on ETH Zurich’s Research Collection (https://doi.org/10.3929/ethz-b-000581447)^22^, and contains 1,312 cells, grouped into 214 Long, 282 Round, 365 Mitotic, 167 Big, and 284 RAR-treated cells.

### Dataset description - eight human immune cell classes

50×50 pixel single-cell multi-channel images were retrieved from ETH Zurich’s Research Collection (https://doi.org/10.3929/ethz-b-000343106)^24^. The dataset consists of a training (89,564 cells) and a test (24,000) dataset with immune cells from multiple healthy donors grouped in eight different classes^12,24^. The eight immune classes consist of CD4+ T cells, CD8+ T cells, and double-negative T cells (CD4- and CD8-), monocytes, dendritic cells (DC), natural killer (NK) cells, B cells, and nucleated immune cells negative for all eight surface markers (see table below, adapted from Severin et al. for clarification^12^). For evaluation, we analysed 8,000 class-balanced cells randomly subsampled from the test dataset spanning across ten healthy donors (n=100 cells per class per donor).

| Marker/staining | Channel (exc. wavelength) | Classes positive for marker |
| --- | --- | --- |
| DAPI | Blue (435-480 nm) | All classes |
| CD3 | Red (650-760 nm) | CD3+ (optionally CD4+ or CD8+) T cells |
| CD4 | Green (500-550 nm) | CD3+CD4+CD8- T cells |
| CD8 | Orange (570-630 nm) | CD3+CD4-CD8+ T cells |
| CD19 | Green (500-550 nm) | CD19+ B cells |
| CD19 | Orange (570-630 nm) | CD19+ B cells |
| CD56 | Orange (570-630 nm) | CD56+ NK cells |
| CD16 | Orange (570-630 nm) | CD16+ NK cells |
| CD14 | Red (650-760 nm) | CD14+ (optionally CD16+ or CD11c+) Monocytes |
| CD11c | Green (500-550 nm) | CD14-CD11c+ Dendritic cells |

### ImageNet DINO-pretrained ViTs

We downloaded the full checkpoint weights (full ckpt) of four different ImageNet-pretrained ViTs architectures (ViT-S/16, ViT-S/8, ViT-B/16, ViT-B/8) published previously (https://github.com/facebookresearch/dino)^18^. We next converted the RGB-pretrained models to be compatible with greyscale multi-channel images. To make the RGB-pretrained models flexible in terms of the number of input channels, we dynamically adjusted the patch embedding. This was accomplished by averaging the RGB-specific embedding weights (torch.nn.conv2d operation in the PatchEmbed module) and then multiplying the resulting average by the number of input channels. Furthermore, we resized the input images from 50×50 pixels (multiplexed immune cells), 100×100 pixels (iPSC quality control data), and 150×150 pixels (iPSC morphologies) to 224×224 pixels and rescaled the pixel intensities to a range between 0 and 1. Compared to the published version, the greyscale images were not normalised with the precomputed RGB-based mean and standard deviations, since the respective values are not compatible with microscopy-based pixel intensities.

### scDINO training

For scDINO we DINO-trained a student and teacher ViT-S/16 architecture (384-dimensional ViT with patch size 16) across all 89,564 cells in the immune cell training dataset, while using all five channels as input data. The single-cell images were resized to 224×224 pixels, resulting in final image dimensions of 224×224×5, and all pixel intensities rescaled to a range between 0 and 1 for each channel. We then adapted the augmentations used in DINO, originally designed for RGB images, for individual greyscale images where possible. These include: horizontal flip, recropping, normalisation, and gaussian blur. We replaced the RGB-specific colorjitter augmentation by using solarisation and brightness adjustments. Then, after augmentations, we normalised each image channel-wise according to the channel-specific mean and standard deviations precomputed on the immune cell training dataset. scDINO was subsequently trained with a batch size of 30 across 100 epochs on 8 GPUs using ETH Zurich’s computational infrastructure. All other hyperparameters, in line with the Vanilla DINO training described previously^18^, and the resulting training log files can be found in the Source Data table of Fig. 2.

### k-NN classification and visualisation

For performance assessment, we evaluated the high-dimensional CLS tokens for each model and channel combination of each cell of the respective dataset using a simple k-NN approach as previously described ^18,26^. In short, we split the respective datasets in a training (80%) and test dataset (20%). Then, we computed and stored the features of the training data. To classify a cell, we next computed the CLS token, and compared it against all previously stored features. The top class was predicted using the top k (k=20) nearest neighbours via weighted voting. If the top-weighted class corresponded to the actual class, the classification was considered correct (known as top-1 k-NN accuracy). The respective k-NN top-1 accuracies were visualised as heatmaps using the ComplexHeatmap^27^ package in R (version 4.0.2).

### UMAP visualisations

UMAPs were generated based on the CLS tokens of the respective model and channel combinations outlined in the figures. For the UMAP calculations, the following parameters were used for the different datasets: iPSC quality control data (distance metric: euclidean, number of neighbours: 30, minimal distance: 0.4, spread: 1.1, number of epochs: 500), iPSC morphologies (distance metric: euclidean, number of neighbours: 20, minimal distance: 0.1, spread: 1, number of epochs: 500), eight immune cell classes (distance metric: euclidean, number of neighbours: 50, minimal distance: 0.3, spread: 1, number of epochs: 500).

## Acknowledgements

We acknowledge the Blutspende Zurich and their anonymous donors for providing PBMCs, the ETH Informatik Dienste for providing compute and storage infrastructure, as well as Yannik Severin and Lucie Kralickova for support, and all members of the Snijder Lab for critical discussions. We gratefully acknowledge funding from the Swiss National Science Foundation (PP00P3_163961 and PP00P3_194809) and the ETH Zurich (ETH-28 20-1).

## Author Contributions

**RP:** Conceptualisation, Methodology, Validation, Formal analysis, Investigation, Resources, Data Curation, Writing - Original Draft, Writing - Review & Editing, Visualisation

**JH:** Methodology, Software, Validation, Formal analysis, Investigation, Resources, Writing - Review & Editing, Visualisation

**SL:** Methodology, Investigation, Resources, Writing - Review & Editing, Funding acquisition

**BS:** Conceptualisation, Methodology, Validation, Investigation, Resources, Writing - Original Draft, Writing - Review & Editing, Visualisation, Supervision, Project administration, Funding acquisition

